# Expansion microscopy reveals subdomains in *C. elegans* germ granules

**DOI:** 10.1101/2022.05.29.493872

**Authors:** Kin M. Suen, Thomas M. D. Sheard, Chi-Chuan Lin, Dovile Milonaityte, Izzy Jayasinghe, John E. Ladbury

**Affiliations:** School of Molecular and Cellular Biology, University of Leeds, UK, LS2 9JT; School of Biosciences, University of Sheffield, UK, S10 2TN

## Abstract

Many membraneless organelles (MLOs) have been shown to form via liquid-liquid phase separation (LLPS). Light and electron microscopy techniques have been indispensable in the identification and characterization of LLPS MLOs. However, for complex MLOs such as the perinuclear germ granule in *C. elegans*, our understanding of how the organelle as a whole is regulated is hampered by 1) the technical limitations in confocal fluorescence imaging in which only a few granule protein markers can be examined at a time and 2) the inaccessibility of electron microscopy. In this study, we take advantage of the newly developed super-resolution method of expansion microscopy (ExM) and in-situ staining of the whole proteome to examine the *C. elegans* germ granule, the P granule. We show that in small RNA pathway mutants, the P granule is smaller compared with wild type animals. Furthermore, we investigate the relationship between the P granule and two other germ granules, mutator foci and Z granule, and show that they are located within the same protein-dense regions while occupying distinct subdomains within this ultrastructure. The experimental workflow developed here will serve as an important tool in our understanding of germ granule biology as well as the biological role of LLPS.

## Introduction

Germ granules, or nuage, are conserved features throughout the animal kingdom. Composed of conglomerates of ribonucleoprotein complexes, germ granules accommodate proteins and RNAs that play essential roles in epigenetics (Voronina et al., 2011). The P granule is a germ granule in *C. elegans* and a number of Argonaute proteins responsible for small interfering RNA (siRNA) pathways are found in this organelle (Seydoux, 2018; Sundby et al., 2021). For example, the piRNA pathway is a small non-coding RNA pathway that plays an important role in the regulation of transposable elements (Czech et al., 2018; Malone and Hannon, 2009; Weick and Miska, 2014) and the Argonaute protein PRG-1 (Piwi Related Gene-1), which binds piRNAs, is localized to the P granule (Batista et al., 2008). Furthermore, disruption of P granules in embryos lead to misregulation of endogenous siRNA (endo-siRNA; Dodson and Kennedy, 2019; Lev et al., 2019; Ouyang et al., 2019). Hence, the P granule is important for small RNA-based epigenetics.

The P granule was the first MLO found to be formed by liquid-liquid phase separation (LLPS; Brangwynne et al., 2009). LLPS is a biophysical process by which biological molecules condense out of the bulk intracellular milieu to form distinct phases, allowing segregation of biological contents without the use of delimiting membranes (Banani et al., 2017; Wheeler and Hyman, 2018). While the molecular rules that govern the formation of LLPS organelles have been intensively studied in the past decade, the implications, or the physiological roles, of LLPS in biological functions are difficult to assess. Further investigation of the spatial arrangements for biomolecules within phase separated MLOs can shed light on the role of LLPS in biological functions. For example, differences in translational activity between the periphery and the core of the P-bodies have been also observed in Drosophila oocytes (Weil et al., 2012); surface tension was found to be responsible for the generation of the internal compartments within the nucleolus to facilitate ribosome biogenesis (Feric et al., 2016).

In this regard, the P granule presents particular challenges in understanding how LLPS facilitates its functions due to the complexity in its composition: more than 40 proteins have been identified as associated with this organelle (Suen et al., 2020; Sundby et al., 2021; Updike and Strome, 2010) and most RNAs in the germline are thought to at least associate with it transiently (Schisa et al., 2001; Sheth et al., 2010). Whilst certain proteins, e.g. PGL-1, are used as markers to visualize P granules by proxy using fluorescence confocal microscopy, how certain treatments or mutations affect the P granule as a whole, or indeed whether sub-variants of P granules exist, is difficult to appreciate relying on light microscopy. Furthermore, the lack of delimiting membrane makes it difficult to ascertain where the boundary of the organelle lies. In contrast, electron microscopy (EM) analysis of *C. elegans* germline tissue has played a critical role in characterizing the P granule without the bias use of specific protein markers. For example, Sheth et al. showed that P granules are not homogeneous throughout but in fact consists of ultrastructures of a crest and a base and that the shape of P granules are dynamic (Sheth et al., 2010). However, the specific proteins that make up these ultrastructures are difficult to identify. While methods such as immunogold staining and more recently correlative light and electron microscopy (CLEM) address this challenge (de Boer et al., 2015; Pacy, 1990), EM-based methods remain accessible only to specialised groups. This makes linking spatial organisations and physiological functions of the P granule difficult.

Given the technical aforementioned limitations of light and electron microscopy, and that protein arrangement within the P granule appear to be organised in the nanometer scale (Putnam et al., 2019; Suen et al., 2020; Wan et al., 2018), we used the advanced super-resolution technique protein retention expansion microscopy (proExM) coupled with indiscriminate proteome stains to visualise the P granule. In proExM, proteins are chemically anchored to a swellable gel which forms a mold of the fluorescently labeled sample. Proteins are non-specifically digested leaving a molecular-scale imprint of the staining covalently linked to the gel. Hydration of the gel allows gel to expand isotropically, leading to the physical separation of the fluorophore and hence an increase in resolution in confocal imaging (Chen et al., 2015; Tillberg et al., 2016). Building on proExM, a whole-proteome staining method (i.e. a pan-protein stain) akin to densitometric stains used in electron microscopy (EM) was established whereby protease digestion of the gel-anchored samples results in the decrowding of intracellular space while providing extra free amines for NHS ester fluorescent-labeling (M’Saad and Bewersdorf, 2020). This results in a protein density-encoded super-resolution images that highlight cellular features such as plasma cell membrane, organelles and spatial heterogeneities of intracellular protein organisation, not restricted by the specificity of antibodies or targeted fluorescent probes.

To improve upon the resolution achievable with standard proExM in conjunction with confocal microscopy, we used the resolution-doubling Airyscan protocol (called Enhanced Expansion Microscopy, EExM; (Sheard and Jayasinghe, 2021)) to image the proExM gels containing pan-protein staining of the bulk proteome in dissected *C. elegans* germlines. Among other intracellular features, we observed P granules at nanoscale resolution (∼40nm in plane and ∼100nm axial). We used this EExM-pan stain pipeline to address two important aspects of P granule biology: the effects of small RNA pathway mutations on P granules and the connectivity between P granules and two perinuclear ribonucleoprotein (RNP) granules, the Mutator foci and the Z granule.

## Results

Immunostaining of fluorescent proteins and pan-protein staining has recently been successfully carried out on whole mount *C. elegans* samples (Yu et al., 2020). However, perinuclear density in the germline that might be P granules were not observed (Yu et al., 2020).We suspected this was due to the presence of the cuticle, and/or the fact that the germline was buried inside the body. proExM and EExM have been successfully performed on tissues without cuticles, as is the case for pan-protein staining for the bulk proteome. Given our interest is in the germline, we decided to test if the current pro/EExM and pan-protein stain protocols are suitable for *C. elegans* germlines extruded by dissection, which would manually remove the cuticle from the tissue-of-interest. Briefly, we dissected 1 day-old adults and used the freeze-crack method to immobilise germline tissues on a coverslip throughout the immunostaining process. Immuno-stained germlines were then anchored to the swellable gel and digested. Prior to gel expansion, it was pan-protein stained with an NHS ester dye to label the bulk proteome. The gel was finally expanded and imaged using the super-resolution mode on a confocal microscope (Fig. 1a).

**Figure 1.**
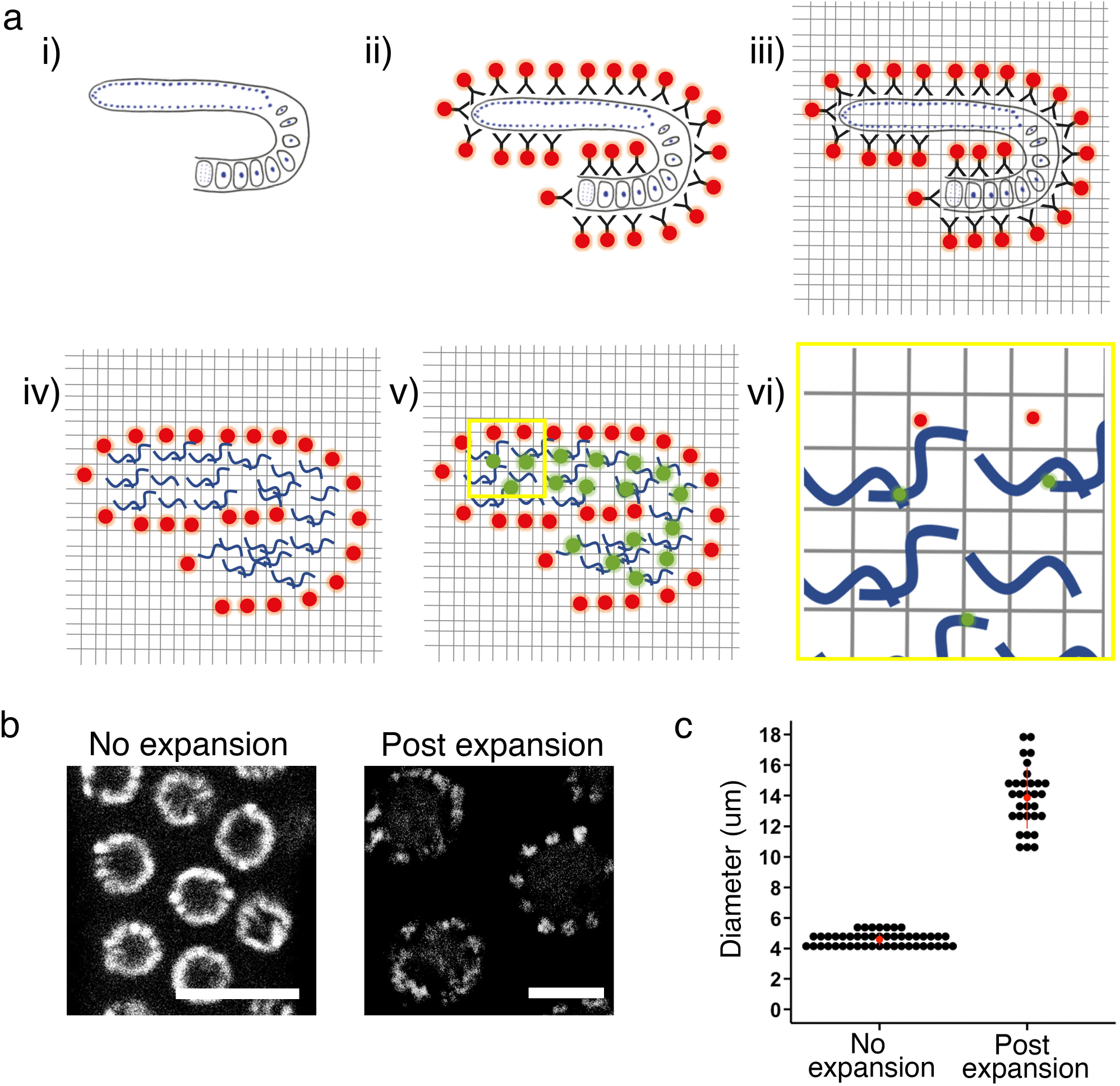
Workflow of expansion microscopy on *C. elegans* germline. a) 3x EExM of immunostained *C. elegans* germline with pan-protein stain. i)-ii) Dissected *C. elegans* germline are stained with primary antibodies followed by fluorescently labelled secondary antibodies. iii) Tissue is then chemically cross-linked to hydrogel which forms a mold of the tissue. iv)Proteins within the tissue are digested prior to v) pan-protein staining of proteins whereby free amines are labelled with fluorescent dye. vi)The hydrogel is expanded 3x and imaged. Resulting resolutions are xy ∼40 nm and z ∼100 nm. b) DAPI staining of the pachytene nuclei in *C. elegans* germline pre-and post-expansion. Scale bar = 10 μm. c) Nucleus diameter measured via DAPI staining to determine expansion factor. The average expansion factor across experiments is 3x. 31 expanded nuclei were measured from 6 independent experiments and 47 non-expanded nuclei were measured from 3 independent experiments.

To determine the expansion factor achieved on the *C. elegans* germline tissue, we measured the diameter of the nucleus in the pachytene region via DNA staining using DAPI. We found that without expansion the nucleus diameter was 4.6 μm ± 0.4 μm (mean ± standard deviation) whereas post-expansion nuclei were measured at 13.9 μm ± 2.0 μm. Hence, the expansion factor is ∼3-fold on average, resulting in a resolution of ∼40nm in plane and ∼100nm axial.

To aid the identification of P granules, experiments were first performed on animals expressing GFP-tagged DEPS-1 (<u>DE</u>fective <u>P</u> granules and <u>S</u>terile-1). DEPS-1 is a scaffold protein essential for P granule formation (Spike et al., 2008). In the pachytene region, pan-protein staining of the *C. elegans* germline proteome reveals a number of intracellular features, including the plasma membrane, nucleus and intranuclear structures (Fig. 2a and video 1). Perinuclear protein-dense structures can be seen using pan-protein staining which are similar to those observed previously with EM (Schisa et al., 2001; Sheth et al., 2010). Importantly, the colocalisation of DEPS-1 to these perinuclear structures confirms these are indeed P granules (Fig. 2a and b) and that the experimental methodology presented here allows for the visualization of P granules. Using super-resolution confocal microscopy, we previously showed that a number of protein condensates, including DEPS-1, are not homogenously distributed throughout the P granule but organised as small protein clusters (Suen et al., 2020). EExM allows us to resolve DEPS-1 condensates further and confirms our previous observation. We determined the number of granules per nucleus in single optical sections using either DEPS-1 or pan-protein staining (Fig. 2c). In most instances, perinuclear DEPS-1 staining coincides with perinuclear protein-dense structures revealed by pan-protein staining as reflected by the similar average number (2.50 via DEPS-1 staining vs 2.75 via pan-protein staining) of granules observed.

**Figure 2.**
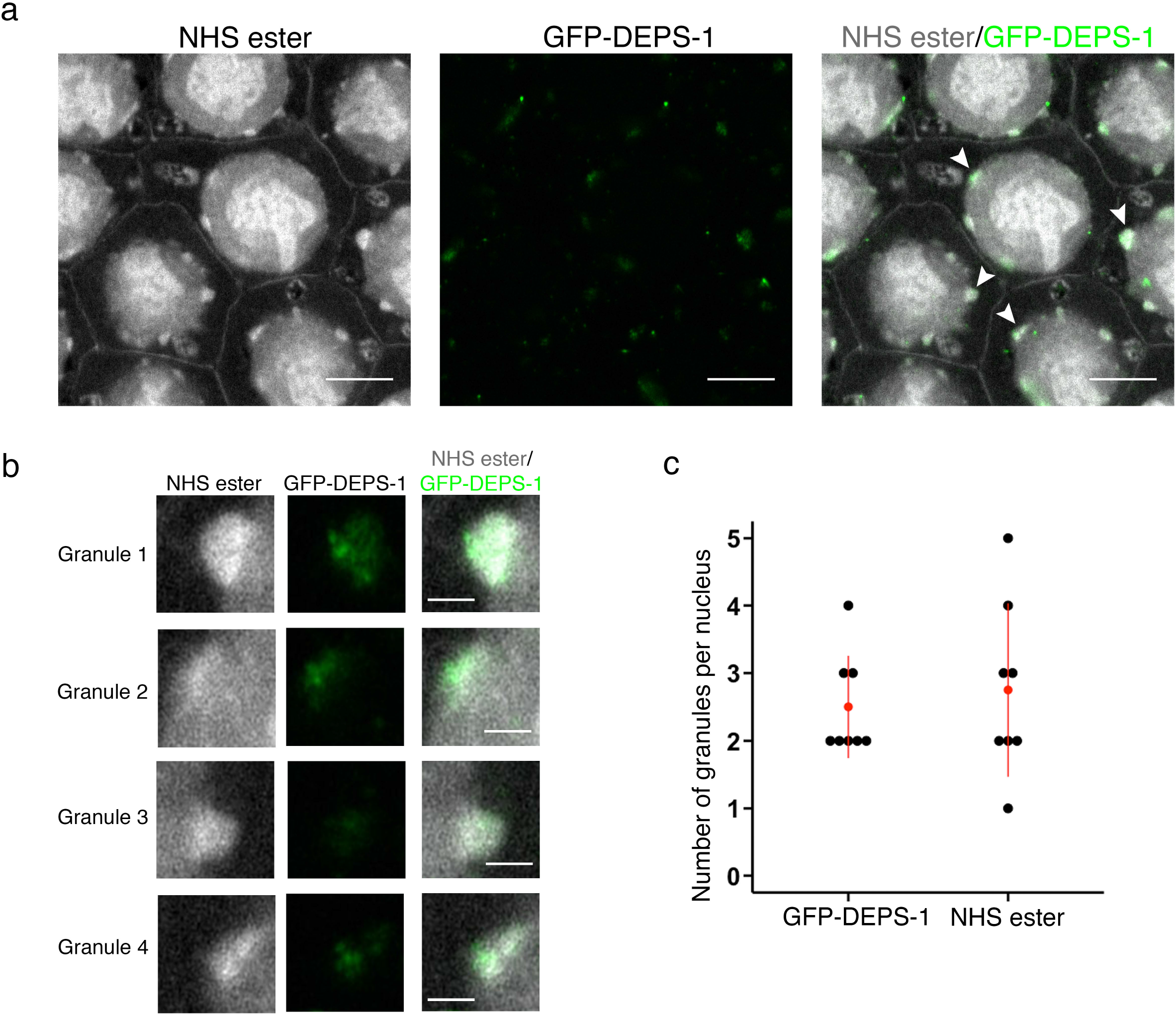
Pan-protein staining reveals P granules as protein-dense perinuclear structures. a) Pan-protein (NHS ester) and anti-GFP staining of animals expressing *gfp::deps-1*. Pan-protein staining reveals a number of features including P granules. GFP-DEPS-1 condensates (green) are localised to the P granule (gray). White arrows highlight P granules that are enlarged in (B). Scale bar = 10 μm. b) Zoomed images of P granules highlighted by arrow heads in (a). DEPS-1 condensates appear as small protein clusters that are localised to P granules. Scale bar = 2 μm. c) Number of granules observed per nucleus in a single optical slice. Granule is defined as perinuclear density observed either via GFP-DEPS-1 staining (green) or pan-protein staining (NHS ester; gray). The average number of granules per nucleus is 2.5 ± 0.8 (GFP-DEPS-1 positive) and 2.8 ± 1.3 (pan-stain positive). 9 nuclei from 3 independent experiments were counted.

### P granule, Mutator foci and Z granule proteins are localised within the same perinuclear protein-dense structures

In addition to the P granule, three other perinuclear germ granules, the Z-granule (Ishidate et al., 2018; Wan et al., 2018), the Mutator foci (Phillips et al., 2012) and the SIMR foci (Manage et al., 2020), have been found in the *C. elegans* germline. Proteins important for various small RNA pathways in the germline often associate with these perinuclear granules. The functions that occur in these different granules/foci are sometimes interconnected. Hence, there has been a suggestion that they are not distinct granules but rather, they are different compartments within the same granule (Sundby et al., 2021). Our pan-stain EExM pipeline is uniquely able to address this question because it allows us to observe perinuclear protein densities non-specifically. We examined proteins that are found in the three different granules: 1) PRG-1 which resides in the P granule, 2) MUT-16 (<u>Mut</u>ator-16), an essential scaffold protein in the Mutator foci (Phillips et al., 2012) and 3) ZNFX-1 (<u>Z</u>inc finger <u>NFX</u>1-type containing homolog) a member of the Z granule (Ishidate et al., 2018; Wan et al., 2018).

We first examined the localization of PRG-1 and MUT-16. We previously showed that PRG-1 and DEPS-1 condensates exist as small protein clusters that intertwine each other (Suen et al., 2020). As expected, PRG-1 is located in perinuclear protein densities (Fig. 3a). We again observed that PRG-1 appears as small protein clusters, and these clusters in general distribute throughout the P granule, defined by the pan-protein staining.

**Figure 3.**
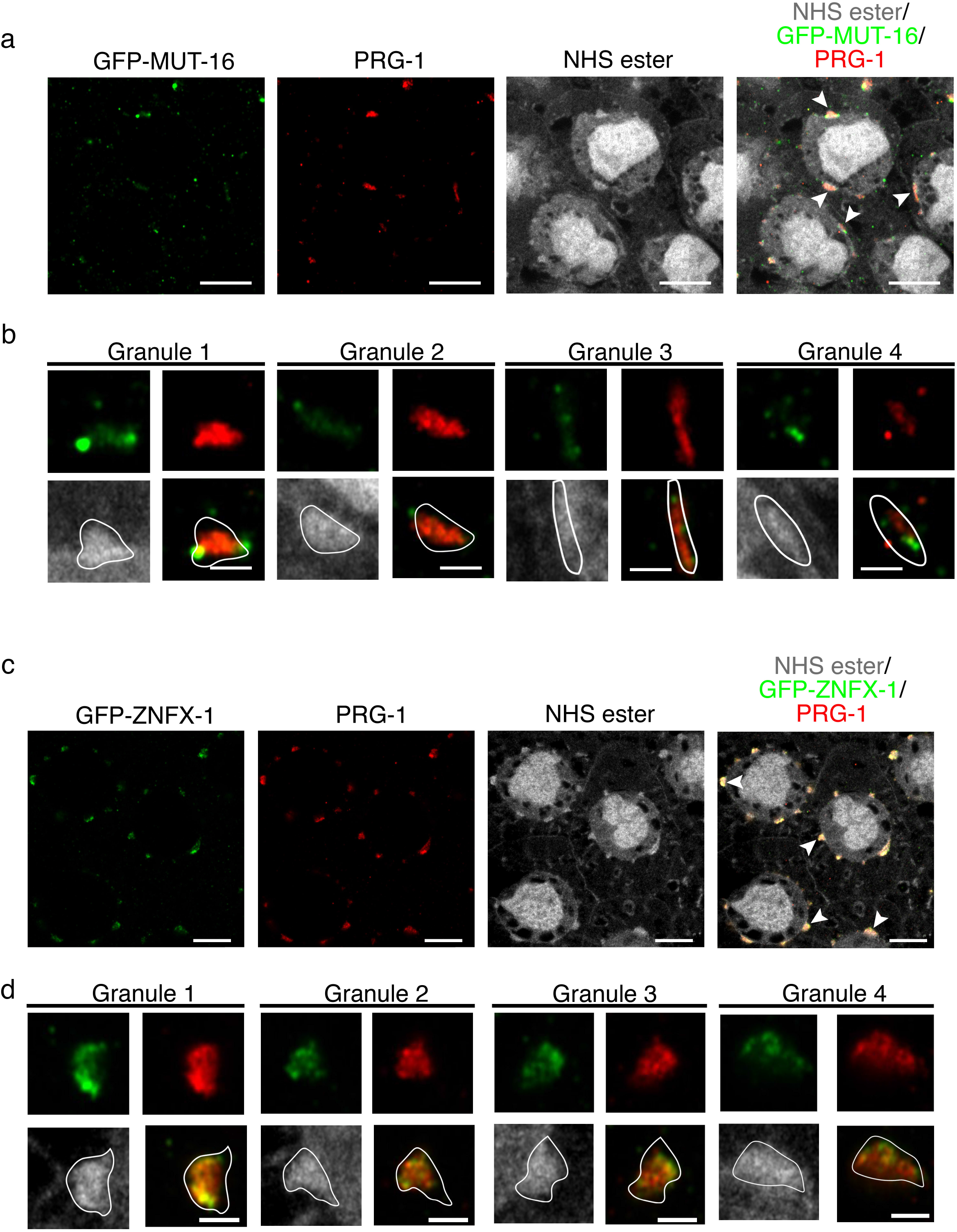
Different small RNA-based epigenetic hubs occupy distinct areas within the same protein-dense structures. a) Pan-protein (NHS ester), anti-PRG-1 and anti-GFP staining of animals expressing GFP-tagged MUT-16. PRG-1 (red) and MUT-16 (green) colocalise to P granules (gray). MUT-16 and PRG-1 occupy distinct areas whereby MUT-16 is frequently observed on the edge of the P granule space. White arrowheads in merged image highlight granules that are enlarged in b). Scale bar = 10 μm. b) Zoomed image of granules marked by arrowheads in (a). PRG-1 (red) exists as small cluster of proteins within the P granule. White line outlines the P granule boundary based on pan-protein staining (gray). MUT-16 (green) appears as single clusters that are either inside the P granule (granules 1, 3 and 4) or outside the P granule (granules 1 and 2). Scale bar = 2 μm. c) Pan-protein (NHS ester), anti-PRG-1 and anti-GFP staining of animals expressing GFP-tagged ZNFX-1. PRG-1 (red) and ZNFX-1 (green) colocalise to P granules (gray). ZNFX-1 and PRG-1 occupy distinct, as well as overlapping areas within the P granule space. White arrowheads in merged image highlight granules that are enlarged in d). Scale bar = 10 μm. d) Zoomed image of granules marked by arrowheads in (c). Both PRG-1 (red) and ZNFX-1 (green) exist as small clusters of proteins within the P granule. White line outlines the P granule boundary based on pan-protein staining (gray). ZNFX-1 is concentrated in areas closer to the cytoplasmic edge of the P granule than PRG-1 (granules 1, 3 and 4). Scale bar = 2 μm.

piRNAs act as the initial recognition factor to identify mRNA transcript. The effectiveness of the piRNAs is dependent on the secondary endo-siRNA pathway mediated by proteins found in the Mutator foci, a MLOs with LLPS characteristics (Phillips et al., 2012; Zhang et al., 2011). MUT-16 is a scaffold protein shown to be essential in the formation of Mutator foci and that loss-of-function mutations in *mut-16* leads to a dramatic reduction in the level of the endo-siRNA 22Gs. It is well-established, via immunostaining for Mutator foci and P granule protein markers, that Mutator foci and P granules are distinct granules. These granules are juxtaposed next to each other when examined using both conventional and super-resolution fluorescent confocal microscopy (Phillips et al., 2012; Wan et al., 2018). We immuno-stained for MUT-16 as a marker for Mutator foci, and PRG-1. In agreement with previous studies, MUT-16 appears as small foci and mostly do not overlap with PRG-1. Interestingly, MUT-16 foci are often located at the periphery of the perinuclear protein dense structures (Fig. 3a). To examine more closely as to whether MUT-16 is within the P granule, we traced the boundary of the perinuclear protein-dense structure of individual granule and overlaid it with the co-staining images of PRG-1 and MUT-16 (Fig. 3b). We found that MUT-16 foci can lie both inside (granules 1, 3 and 4) and outside the P granule boundary (granules 1 and 2).

Together with the Argonaute protein WAGO-4, ZNFX-1 plays an essential role in the transgenerational inheritance of RNAi (Ishidate et al., 2018; Wan et al., 2018). It was shown that ZNFX-1 segregates from the P granule protein PGL-1 at the Z2/Z3 embryonic stage which leads to the formation of Z granules (Wan et al., 2018). We found that ZNFX-1 and PRG-1 exist within the same perinuclear protein densities but occupy different areas, as well as some overlapping positions, indicating that the Z granule do not constitute a separate protein dense structure but in fact are compartments, or subdomains (Fig. 3c; Ishidate et al., 2018), within the P granule. Furthermore, ZNFX-1 frequently occupies the area closer to the cytoplasmic edge than the nuclear membrane edge of the P granule (Fig. 3d).

### Gross changes to P granule morphology in *deps-1* and *mut-16* mutants

*deps-1* was first identified as a gene required for the correct localisation of PGL-1 which is a critical component of P granules. In the adult germline, DEPS-1 is thought to be its most foundational member in the P granule, i.e. the localisation of all other P proteins are dependent on the presence of wild type *deps-1* (Spike et al., 2008). While this hierarchy was identified by examining the localisation of specific P granule proteins, how P granules as an organelle is affected by *deps-1* mutations is unknown. We performed pan-protein staining on *deps-1 (bn124)* null animals to investigate whether the number and morphology of P granules are affected (Fig. 4a and video 2). In *deps-1 (bn124)* null animals, perinuclear densities appear to be flatter overall compared with wild type animals (Fig. 4d). Furthermore, protein densities containing PRG-1 can be infrequently seen as granules almost dissociated from the nuclear membrane (Fig. 4b, supplemental Fig. 1).

**Figure 4.**
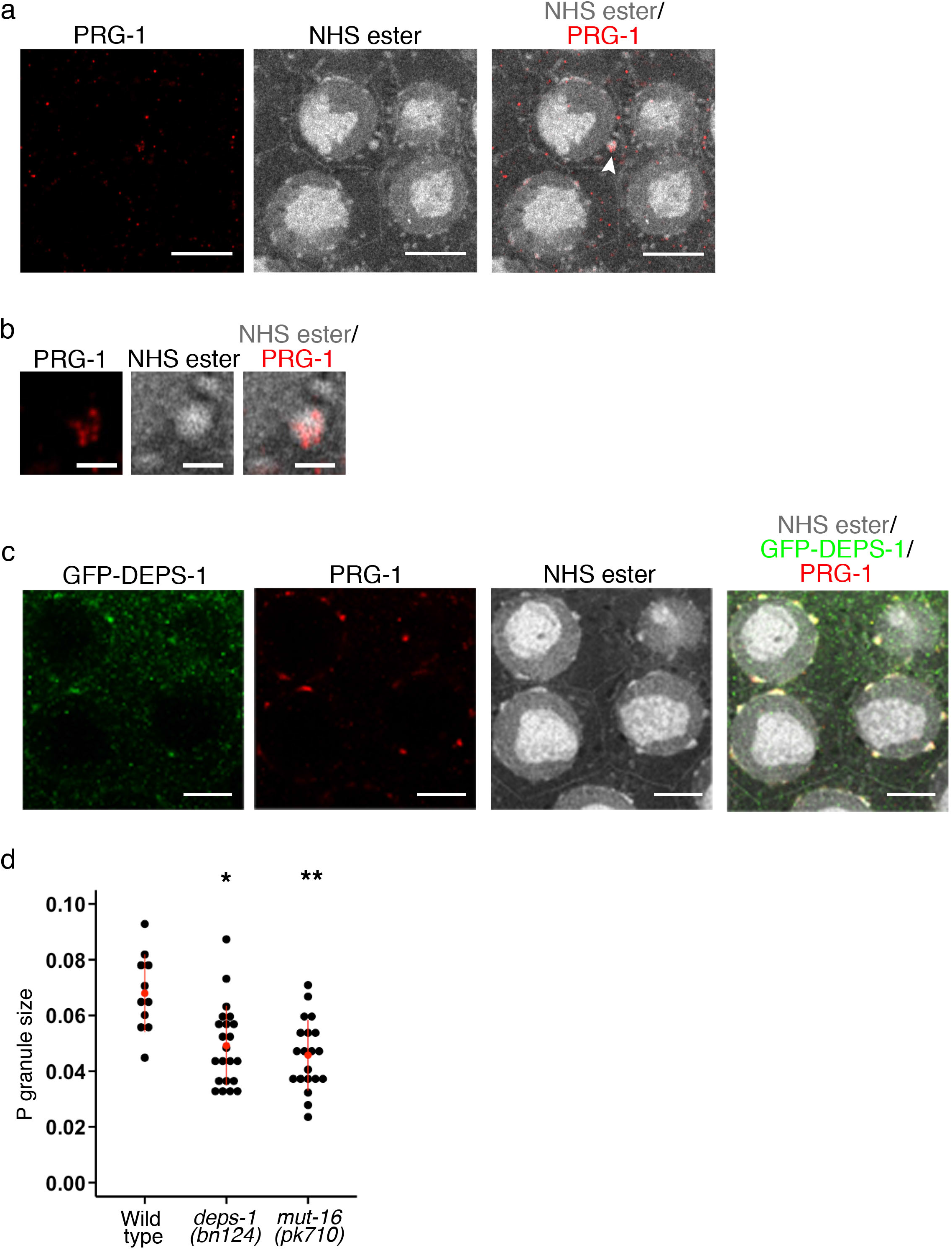
P granules are malformed in animals defective in small RNA pathways. a) Pan-protein (NHS ester) and anti-PRG-1 staining of *deps-1* mutant animals. *deps-1 (bn124)* mutation leads to an overall reduced number of perinuclear protein-dense structures. Scale bar = 10 μm. b) Zoomed image of a P granule (highlighted by arrowhead in (a) containing PRG-1 that is dissociated from the nuclear membrane in a *deps-1 (bn124)* mutant animal. Scale bar = 2 μm. c) Pan-protein (NHS ester), anti-DEPS-1 and anti-PRG-1 staining of *mut-16 (pk710)* mutant animals. *mut-16* mutation leads to a reduction in the size of P granules. Scale bar = 10 μm. d) P granule size in wild-type animals, *deps-1 (bn124)* and *mut-16 (pk710)* mutants. P granule size was calculated by measuring the maximum length of the P granule perpendicular to the nuclear membrane and normalizing it to the diameter of the nucleus. P granules are smaller in both mutants compared with wild-type animals. * p value < 0.001 and ** p value < 0.0001. Video 1 Z Stack images of the pachytene region of *C. elegans* germline in wild-type animals. Scale bar = 10 μm. The germline is pan-protein stained using NHS ester Atto 488. Video 2 Z Stack images of the pachytene region of *C. elegans* germline in *deps-1 (bn124)* mutant animals. Scale bar = 10 μm. The germline is pan-protein stained using NHS ester Atto 488. Video 3 Z Stack images of the pachytene region of *C. elegans* germline in *mut-16 (pk710)* mutant animals. Scale bar = 10 μm. The germline is pan-protein stained using NHS ester Atto 488.

We previously showed that mutations in Mutator foci affect DEPS-1 perinuclear localization. Pan-protein staining reveals that similar to the *deps-1 (bn124)* mutant, *mut-16 (pk710)* mutant animals also exhibit much diminished perinuclear protein densities (Fig. 4c-d and video 3). Given that *deps-1 (bn124)* mutation does not lead to a dramatic reduction in small RNA levels (Suen et al., 2020) but is required for the correct localization of other proteins (Spike et al., 2008; Suen et al., 2020), whereas *mut-16* mutation causes almost the complete loss of 22Gs but not P granule protein localization (Phillips et al., 2012; Suen et al., 2020; Zhang et al., 2011), the morphology of these perinuclear protein densities are dependent on both small RNA and protein levels.

## Discussion

The P granule in *C. elegans* has been fundamental for the discovery that MLOs can be formed by LLPS. It is a site in which the protein components of various small RNA pathways are located. Hence, understanding how small RNA-based epigenetics is facilitated or driven by LLPS in the P granule will shed light on the biological relevance of LLPS.

Both electron and fluorescent light microscopies have played invaluable roles in the characterization of P granules: electron microscopy led to the first observation of P granules as perinuclear densities that are non-membrane bound (Schisa et al., 2001; Sheth et al., 2010) and fluorescent light microscopy has led to the identification of protein and RNA components in these perinuclear germ granules (Sundby et al., 2021; Updike and Strome, 2010). However, identifying the localization of specific proteins among ultra-structures revealed in EM is extremely challenging and EM is limited in its accessibility preventing its wide-spread use, for example, in screening the morphology of P granules in mutants. Light microscopy, whilst it is accessible to non-specialist groups, relies on the immunostaining for specific proteins as markers for specific compartments. In complex MLOs such as the P granule, this ‘biased’ approach does not allow examination of the organelle as a whole. In this work, we adapted pan-protein staining and EExM to overcome these limitations and present a workflow that can be easily applied to examine P granules, among other cellular features, at nanoscale resolution.

We examined the localization of proteins that make up three small RNA pathways: PRG-1 for the piRNA pathway, MUT-16 for secondary endo-siRNAs and ZNFX-1 for transgenerational inheritance of RNAi response. While PRG-1 is a known as a P granule protein due to its colocalization with PGL-1 (Batista et al., 2008), MUT-16 and ZNFX-1 are described as independent foci or granules that do not mix with PGL-1 (Phillips et al., 2012; Wan et al., 2018). Whether or not Mutator foci or Z-granule occupy the perinuclear protein density, or nuage, is unclear. Here we addressed this important question using our pan-protein staining with EExM workflow. We found that Mutator foci can exist both within and outside the perinuclear protein density marked by PRG-1 as P granule, whilst ZNFX-1 always exists within the same perinuclear protein dense structures as PRG-1. This means that ZNFX-1 should be considered as a sub-compartment, or subdomain, of P granule, rather than an independent granule. Furthermore, pan-protein staining provides context to protein localization by defining the border of the P granule. We show that MUT-16 is often observed to be located at the periphery of the P granule close to the nuclear membrane. In contrast, ZNFX-1 can be found frequently concentrated to the cytoplasmic edge of the P granule.

These observations raise two important questions. Firstly, how should P granules, Z granules and Mutator foci be defined? ZNFX-1 and MUT-16 clearly can occupy distinct areas from PRG-1, a P granule protein, in the perinuclear protein dense structure. However, conventionally, the P granule describes the entire perinuclear protein dense structure observed in EM studies (Schisa et al., 2001; Sheth et al., 2010). Many proteins have been classified as ‘P granule’ proteins primarily using confocal microscopy with PGL-1 as a protein marker for P granules. With the advent of super-resolution microscopy it would be worth re-examining the localization of these P granule proteins. This might lead to the identification of other sub-domains within the P granule.

Secondly, since these granules co-exist in the perinuclear protein dense structures, it remains to be determined how Mutator foci, Z granule and P granule proteins segregate into different areas. Feric *et al*. shows that the compartments within the nucleolus are linked to the steps in ribosomal biogenesis and this segregation is driven by the differences in droplet surface tensions (Feric et al., 2016). Given that RNA species, both long and short, are main players in the epigenetic pathways, it would be reasonable to hypothesize that RNA contributes to the dynamic formation of subdomains within the P granule.

The contribution of RNA in P granule structural integrity is supported by the gross morphological changes observed in *mut-16* mutant animals, which exhibits a significant loss in specific small RNA populations (Zhang et al., 2011). Phillips *et al*. previously showed that PGL-1 localisation is not disrupted by mutations in Mutator foci genes, including *mut-16* (Phillips et al., 2012), suggesting that the reduced size in P granules might be driven by the loss of small RNA. In contrast, *deps-1* mutation leads to a loss of PGL-1 from the perinuclear region (Spike et al., 2008) while exhibiting a less significant loss in small RNAs compared with *mut-16* mutants (Suen et al., 2020). Hence, morphology of the P granule is dependent on both proteins and RNA.

Whilst previous EM studies show that the P granule consists of a crest and a base (Sheth et al., 2010), we were not able to observe these features using pan-protein staining. It will be interesting to examine the P granule at a higher resolution through higher orders of EExM, which might allow us to observe these features, however the risks of spatial distortions that accompany the combination of multicellular tissue imaging with 10x or greater EExM need to be carefully managed (Sheard et al., 2021). It is also possible that the crest is composed of other materials e.g. RNA, which would not be stained by the pan-protein stain but could be labelled by nucleic acid-based approach.

Our study has focused on observing subdomains within the P granule and determining the effect of small RNA pathway mutants. However, a number of cellular features are visible under pan-protein staining and it would be important to identify these.

## Materials and Methods

### General animal maintenance

Animals were fed with HB101 and maintained at 20 °C on NGM plates. Strains used in this study are listed in Supplemental table 1.

### Worm dissection and immunostaining

Microscope slides were incubated with 0.01% poly-lysine solution for 1 hr at room temperature and excess liquid was removed by tissues. 1-day-old adults were dissected for germline on the poly-lysine coated slides in 9 μl of 1 mM levamisole (diluted in M9) using 21G needles to remove their heads or tip of the tails. Extruded germlines were then fixed with 4% paraformaldehyde for 10 min at room temperature. After 10 min, the solution was diluted 3-fold, using a gel-loading pipette tip to remove excess liquid (i.e. leaving 9 μl). A rectangular coverslip was then put on top of the dissected samples perpendicular to the slide and placed on top a prechilled metal block (chilled by dry ice) for at least 10 min. The slides were then removed from the metal block and coverslip lifted up in one smooth motion. The slides were placed in prechilled methanol at –20 °C methanol for 20 min. Fixed samples were washed with PBS-T (PBS supplemented with 0.1% tween-20) prior to primary antibody addition. Primary antibodies were incubated with the samples at 4 °C for overnight. Secondary antibodies and DAPI were incubated at 37 °C for 1 hr in the dark. Antibodies used: anti-PRG-1 (Custom, 1:500); anti-mouse GFP (Thermofisher, A-11120; 1:500). Anti-rabbit Alexa fluor 594 secondary antibody (Thermofisher A11012) was used at 1:500 and anti-mouse Atto 647 secondary antibody (Sigma, 50185) was used at 1:160.

### Gel expansion

Dissected, immunostained germlines mounted on microscope slides were washed twice PBST and then twice with PBS. The germlines were incubated with 0.1 mg/ml Acryloyl-X SE (stocks stored as 10 mg/ml in DMSO and diluted to working concentration in PBS; Thermo Fisher, A20770) overnight at 4 °C. Slides were washed three times with PBS next afternoon. Excess liquid was removed as much as possible without drying the germlines prior to the addition of monomer solution (8.6 mg/ml sodium acrylate (Fluorochem, 461652), 2.5 mg/ml acrylamide, 0.15 mg/ml N,N’-Methylenebisacrylamide (Merck, M7279), 11.7 mg/ml NaCl in 1x PBS). Samples were incubated in monomer solution for 30 min at 4 °C. Monomer solution was then removed as much as possible. 100 μl of polymer solution (94 μl monomer solution, 4 μl 1x PBS, 1 μl 10 % APS and 1 μl 10 % TEMED) was added to the sample and a coverslip was gently placed on top of the sample. Gelation was performed in a humid chamber at 37 °C for 2 hrs. Coverslip was carefully removed using a razor blade. The gel was detached from the slide using a razor blade and placed into a small petri dish. Digestion buffer (50 mM Tris pH 8.0, 1 mM EDTA, 0.5% Triton X-100, 0.8 M guanidine HCl and 1: 100 Proteinase K) was added in sufficient amount to cover the gel to digest the germline and the gel was digested overnight at room temperature. Digestion solution was removed the next morning and the gel was washed with PBS twice, transferred to a new petri dish and washed with 100 mM sodium bicarbonate twice. The gel was then incubated in 10 μM NHS ester Atto 488 in 100 mM sodium bicarbonate with gentle rocking at room temperature for 1.5 hrs. The gel was washed with PBS once and transferred to a larger petri dish (e.g. 10 cm petri dish). Deionised water was added to the dish 3-5 times, 30 – 60 min each until expansion reaches plateau.

### Confocal microscopy

50 mm μ-dishes (Ibidi, 81131) were coated with 01% poly-lysine to prevent drift during imaging. Images were taken on a Zeiss LSM880 inverted microscope using the Airyscan mode.

### P granule size analysis

The maximum height of the P granule perpendicular to the nuclear membrane was measured. To account for the expansion factor in the individual samples, the height of the P granule is divided by the diameter of the nucleus.

## Supporting information

Supplemental table 1, animals strains used in this study and cited in Materials and Methods

Video 3, mut-16 mutant animal

Video 2, deps-1 mutant animal

Video 1, wild type animal

## Legends

**Supplemental Figure 1.**
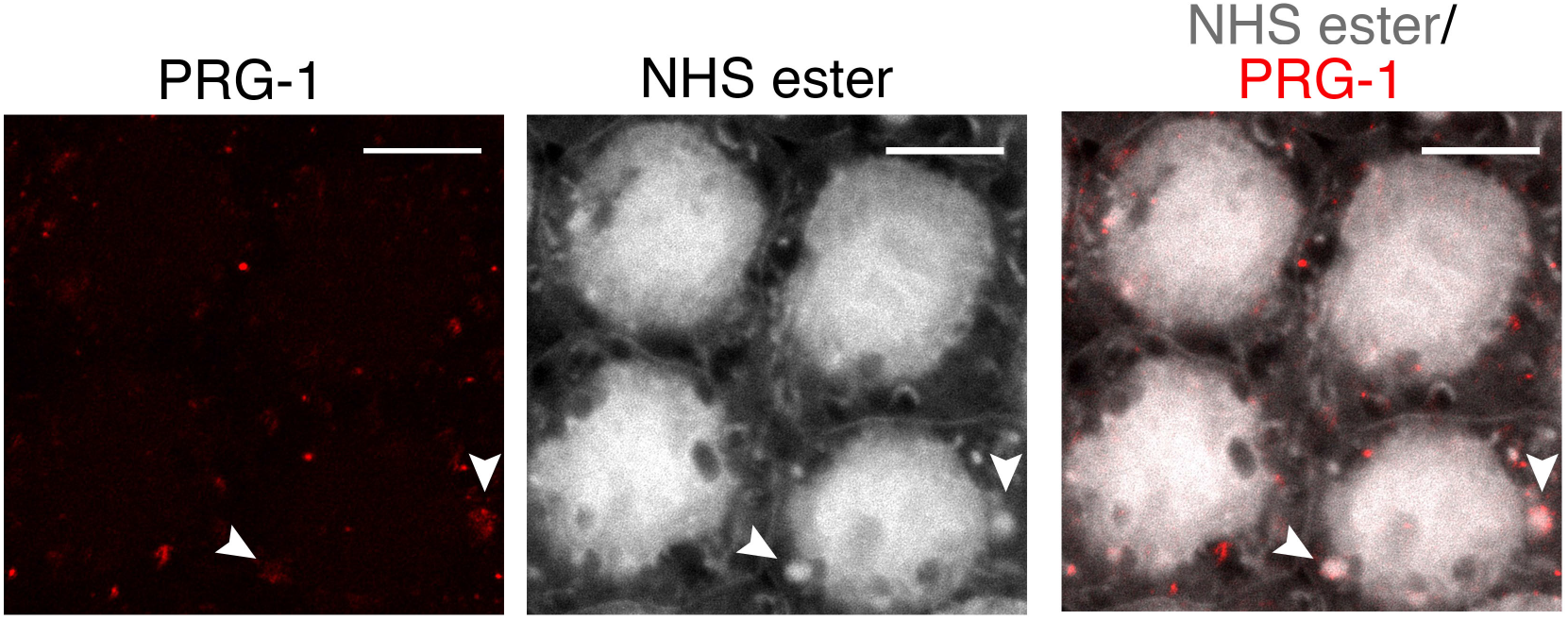
P granules in *deps-1* mutants dissociates from nuclear membrane. Pan-protein and anti-PRG-1 staining of *deps-1 (bn124)* mutant. Arrow heads highlight 2 P granules that are mislocalised. Scale bar = 10 μm.

