## Supplementary figures and images for "Expansion microscopy reveals subdomains in *C. elegans* germ granules"

### Video 1, wild type animal

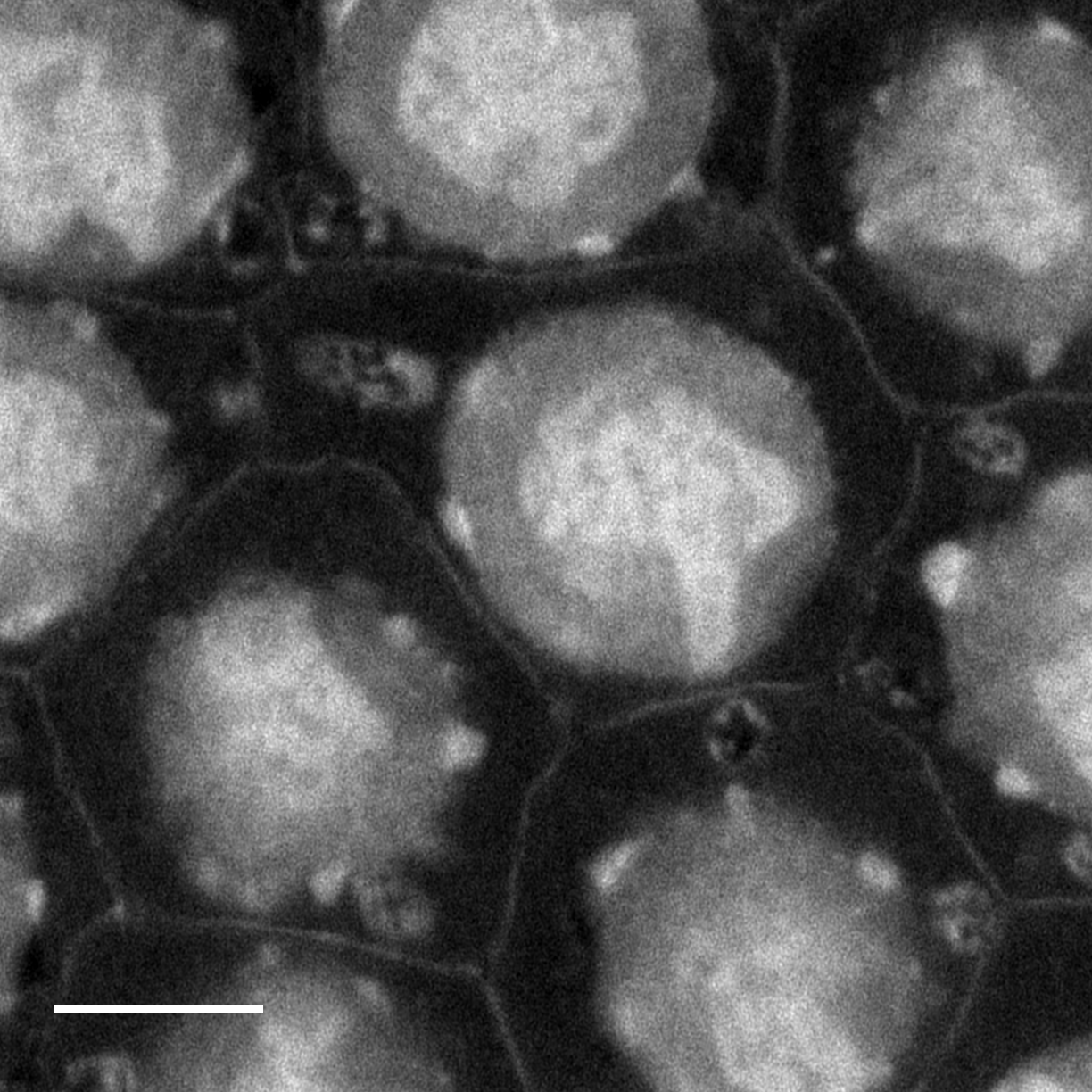
